# Mammalian Cells Engineered to Produce Novel Steroids

**DOI:** 10.1101/261362

**Authors:** Emma S. Spady, Thomas P. Wyche, Nathanael J. Rollins, Jon Clardy, Jeffrey C. Way, Pamela A. Silver

## Abstract

Steroids can be difficult to modify via traditional organic synthesis methods, but many enzymes regio- and stereo-selectively process a wide variety of steroid substrates. We tested whether steroid-modifying enzymes could make novel steroids from non-native substrates. Numerous genes encoding steroid-modifying enzymes, including some bacterial enzymes, were expressed in mammalian cells by transient transfection and found to be active. We made three unusual steroids by expression in HEK293 cells of the 7α-hydroxylase CYP7B1, which was selected because of high native product yield. These cells made 7α,17α-dihydroxypregnenolone and 7β,17α-dihydroxypregnenolone from 17α-hydroxypregnenolone, and produced 11α,16α-dihydroxyprogesterone from 16α-hydroxyprogesterone. The latter two products resulted from previously unobserved CYP7B1 hydroxylation sites. A Rosetta docking model of CYP7B1 suggested that these substrates’ D-ring hydroxylations may prevent them from binding in the same way as the native substrate, bringing different carbons near the active ferryl oxygen. This new approach could use other enzymes and substrates to produce many novel steroids for drug candidate testing.

## Introduction

Steroids represent an important class of therapeutically beneficial molecules, but have significant side effects that limit their use. Steroid hormone analogs take advantage of the wide-ranging effects of their endogenous counterparts, such as immunosuppression and increased bone density, and thus are used to treat conditions ranging from arthritis to anemia. Unfortunately, the global responses steroids alter result in side effects, such as diabetes and osteoporosis for glucocorticoids, or masculinizing effects for androgens^1–3^. They are active at bloodstream concentrations in the nanomolar range and readily cross cell membranes^4,5^. While steroid hormone receptors have known activators and repressors, new combinations of modifications around the central steroid scaffold could improve binding affinity, pharmacokinetic properties, and receptor specificity^6^.

Steroids must be modified stereo- and regio-specifically to result in useful drugs. Steroid receptor binding pockets are sensitive to minor modifications; a single hydroxylation or methylation can cause a steroid to bind a different receptor, dramatically changing its biological effects^6^. Due to its lack of activated carbons, the steroid ring structure is difficult to modify while preserving its integrity^7^. Traditional chemical processes for steroid modifications are elaborate and frequently less selective than their biological counterparts, to the extent that whole, un-engineered fungi are used as industrial biocatalysts^7–9^.

Metabolic engineering can provide an alternative source of novel steroid therapeutics. The enzymes involved in steroid hormone synthesis modify a large variety of substrates, but still perform regio- and stereoselective reactions^10^. These enzymes could accept a wide variety of synthetic steroid substrates and predictably modify them to create novel products. A single enzymatic step could replace a long organic synthesis, and thus allow for extensive exploration of the steroid drug biochemical space. Exogenous expression of steroid enzymes has already been used to produce complicated steroids intracellularly in yeast and mammalian cells, with glucocorticoids being of particular interest^11,12^. However, intentional biosynthesis of chemically novel steroid products using metabolically engineered mammalian cells has not to our knowledge been attempted.

Here we report the synthesis of novel steroids by expressing various steroid-modifying enzymes in mammalian cells and providing them with non-native substrates. We isolated three new steroids by expressing the steroid 7α-hydroxylase CYP7B1, which had the highest yield of the mammalian enzymes tested. This approach could be used to synthesize a wide variety of steroids for which traditional organic synthesis is simply not feasible.

## Results & Discussion

Our strategy for novel steroid synthesis is to express genes encoding steroid-modifying enzymes in mammalian cells, and then supply the cells with non-native substrates that will result in novel products (Figure 1). To identify candidates, we expressed a panel of enzymes in mammalian cells and confirmed their function on native substrates. We then selected one of the best-performing enzymes, CYP7B1, and chose non-native substrates that would result in novel products, assuming that the enzyme modifies the substrates selectively. We isolated the products from the enzyme acting the alternative substrate, and confirmed the steroids’ structures via NMR. Finally, we explained how the observed unexpected products may have arisen with a Rosetta docking model for the substrates in the enzyme’s binding pocket.

**Figure 1.**
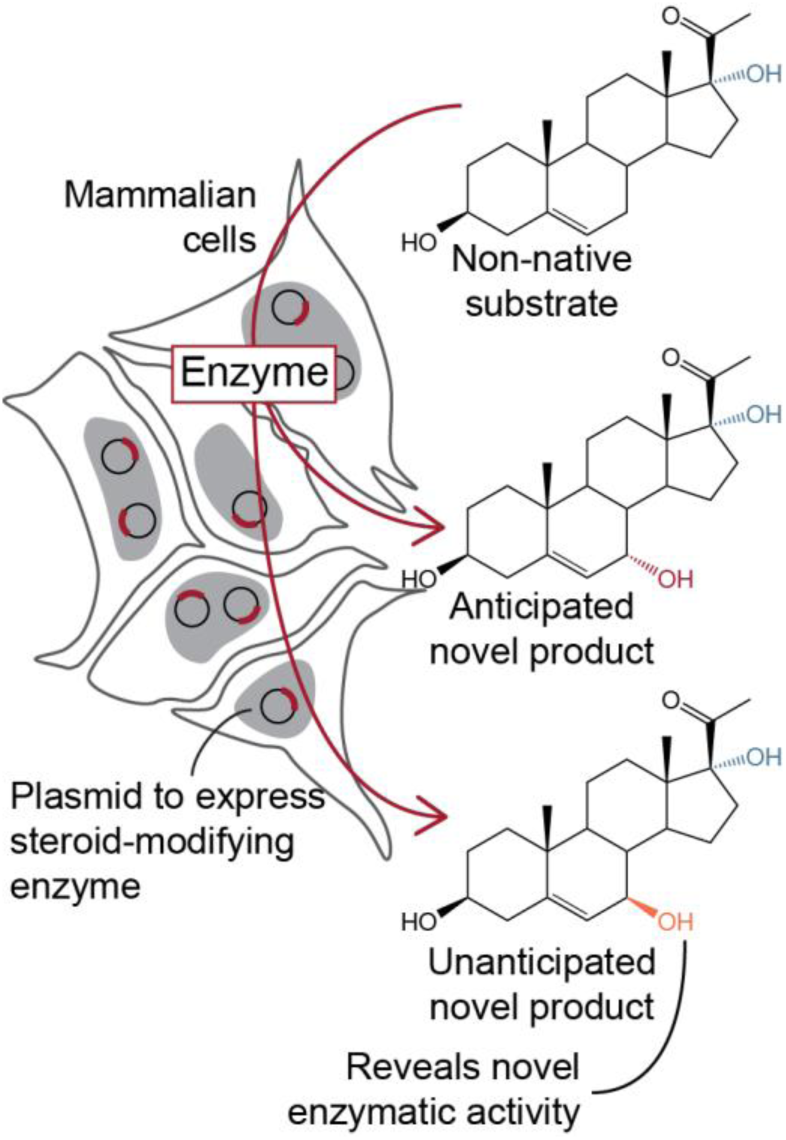
Overview of our method for novel steroid biosynthesis. Steroid-modifying enzymes are expressed heterologously in mammalian cells and exposed to a non-native substrate, resulting in novel steroid product(s).

Seventeen steroid-modifying enzymes were selected for expression in the human cell line HEK293. The enzymes were identified based on their gene sequence availability and the diverse reactions they catalyze (Table S1). Most of these enzymes are cytochrome P450s, and many are hydroxylases; the group also contains dehydrogenases, a methyltransferase, and a Baeyer-Villiger monooxygenase. Mammalian steroid-modifying enzymes are generally associated with the mitochondria or endoplasmic reticulum and may have signal sequences targeting them to these organelles; these sequences were retained. Bacterial enzymes were expressed without signal sequences and presumably localized to the cell cytoplasm. The HEK293 cell line was chosen because of its hardiness and ease of transfection, and because most of the enzymes tested were from humans. Use of human cells also enabled the enzymes to access cholesterol when required as a substrate.

Expression constructs encoding each enzyme were transfected into HEK293 cells, and activity of each enzyme was tested on a native substrate and assayed via LC-MS. Negative enzyme activity controls consisted of transfecting the same plasmid backbone, but with a fluorescent reporter protein under the constitutive promoter. Cells were incubated for two days with a 40 μM solution of a native substrate for the relevant enzyme. The substrate concentration was chosen because of the low solubility of the less-hydroxylated steroids in media^13^. Lipids were extracted from the spent cells and media and measured by LC-MS^14,15^. The yield per milliliter of spent media was determined for each product when a standard was available.

Of the enzymes tested, the human 7α-hydroxylase CYP7B1 and the *Pseudomonas* cyclopentadecanone monooxygenase CpdB had the highest yields. The former produced 1.4 μg of 7α-hydroxypregnenolone per mL (**2**), while the latter produced 2.6 μg of testololactone per mL (**10**) (Figure 2A, Figure S1). Eight other steroid-modifying enzyme plasmids resulted in significantly more product than the fluorescent protein control when transfected (Table S1). These included two more bacterial enzymes, Δ^1^-KSTD2 and CYP154C3 (Figure S1). CYP7B1 was pursued further, as it is a well-studied human enzyme^16,17^. The high 7α-pregnenolone (**2**) yield implied CYP7B1 had a good chance of processing alternative substrates into detectable quantities of novel steroids.

**Figure 2.**
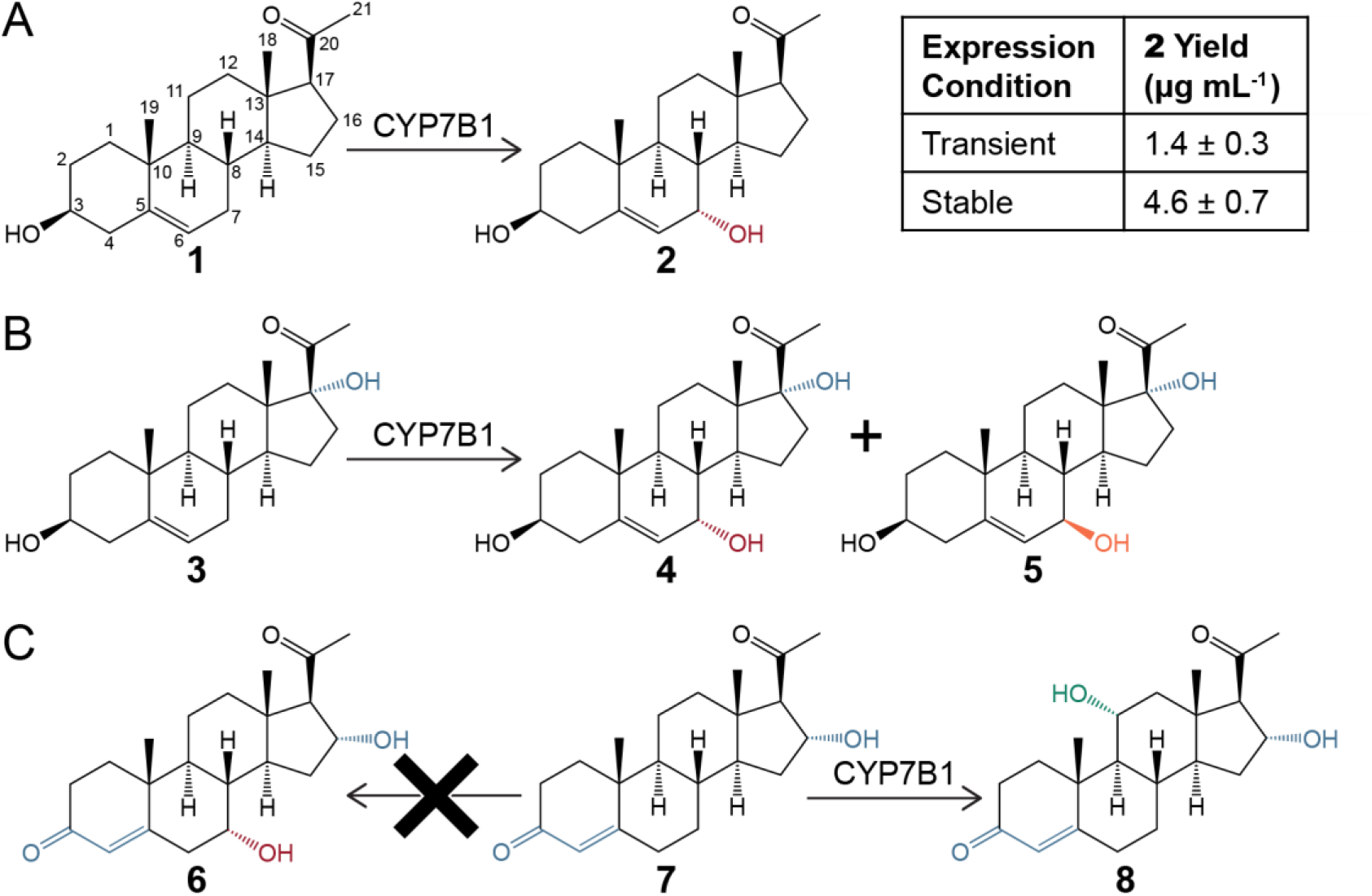
Steroid-modifying reactions observed from CYP7B1. A) CYP7B1 hydroxylates native substrate pregnenolone (**1**) to form 7α-hydroxypregnenolone (**2**). Yield is in μg of steroid per mL of media extracted. B) CYP7B1 hydroxylates non-native substrate 17α-hydroxypregnenolone (**3**) to form novel product isomers 7α,17α-dihydroxypregnenolone **(4)** and 7β,17α-dihydroxypregnenolone (**5**). C) CYP7B1 hydroxylates non-native substrate 16α-hydroxyprogesterone (**7**) to form 11α,16α-dihydroxyprogesterone (**8**). The anticipated product 7α,16α-dihydroxyprogesterone (**6**) was not observed.

The CYP7B1-expressing cells’ yield of **2** improved under different expression conditions. This ensured that the enzyme’s action on alternative substrates would result in sufficient product for structure determination. The CYP7B1 electron transfer partner, cytochrome P450 oxidoreductase (POR), was expressed alongside CYP7B1 to prevent POR from limiting steroid hydroxylation. A modest 34% increase in **2** yield was achieved by transfecting CYP7B1 plasmid into stable POR-expressing HEK293 cells (Figure S2A). A CYP7B1-expressing stable pool was constructed to enable longer exposure of cells to substrate. These cells could be plated at low density and grown to confluency in the same media, allowing for eight days of incubation with substrate instead of 48 hours. HEK293 cells were transfected with the CYP7B1 vector used previously, and were selected on puromycin. This stable pool improved **2** yield by 229%, to 4.6 μg mL^−1^ (Figure S2A). The CYP7B1-expressing stable pool was used in subsequent steroid syntheses, and the POR stable pool was not used.

Stable CYP7B1-expressing cells were incubated with two alternative substrates that we predicted would result in novel products. We anticipated that CYP7B1 would convert 17α-hydroxypregnenolone (**3**) to 7α,17α-dihydroxypregnenolone (**4**) (Figure 2B). Similarly, CYP7B1 was expected to hydroxylate 16α-hydroxyprogesterone (**7**) to produce 7α,16α-dihydroxyprogesterone (**6**) (Figure 2C). The CYP7B1 stable pool cell line was plated at low density and cultivated in the presence of an alternative substrate for eight days. Lipids were extracted and detected by LC-MS as in the native substrate experiments. The lipids were compared to those from identically treated negative control stable pool cells, which expressed GFP instead of an enzyme. The desired hydroxylation product peaks were identified by having the correct mass and only being present with CYP7B1 expression. This separated the desired steroids from any hydroxylated isomers due to spontaneous degradation.

Peaks with masses corresponding to hydroxylation of the alternative substrates were detected exclusively in CYP7B1-expressing cells. CYP7B1 cells provided with alternative substrate **3** yielded two dihydroxypregnenolone peaks not present in the control (Figure 3A). As dihydroxypregnenolones lose water during ESI-MS, the product peaks contained *m/z* = 331.227, 313.217, and 295.206, corresponding to [M+H-H_2_O]^+^, [M+H-2H_2_O]^+^, and [M+H-3H_2_O]^+^ions. The later, high signal molecule **4** eluted at 2.7 minutes, and the earlier, low signal molecule **5** eluted at 2.3 minutes. 0.7 mg of **4** and 0.4 mg of **5** were subsequently purified, implying a yield of 1.6 μg mL^−1^ and 0.91 μg mL^−1^, respectively.

**Figure 3.**
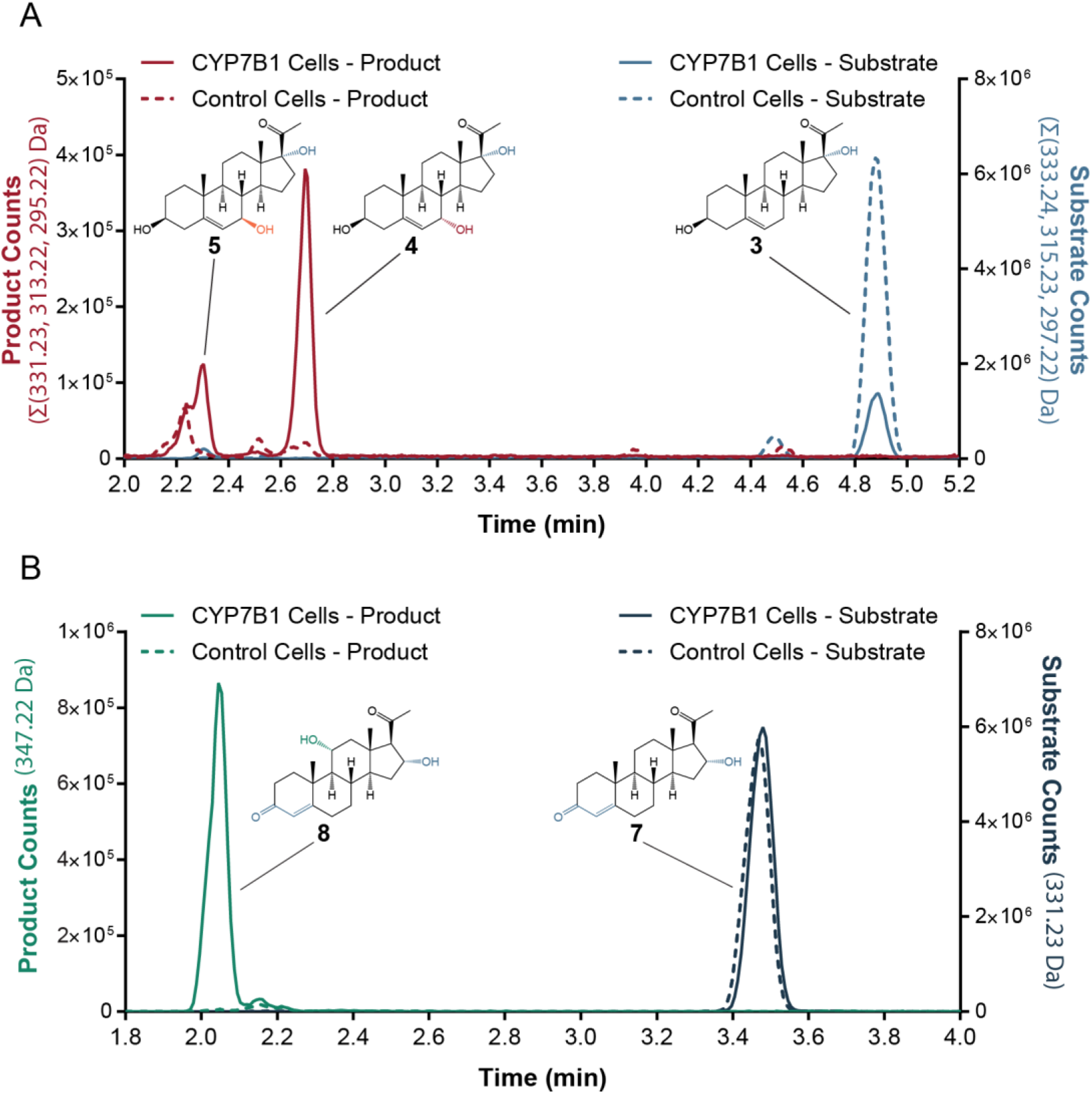
Extracted ion mass chromatograms comparing products of cells supplied with **3** and **7**. CYP7B1 cells stably expressed that enzyme, while the negative control cells stably expressed green fluorescent protein. CYP7B1 cell samples were diluted tenfold. A) Only CYP7B1 cells exposed to **3** produced peaks corresponding to the [M+H-H_2_O]+, [M+H-2H_2_O]+, and [M+H-3H_2_O]+ ions of **4** and **5**. B) Only CYP7B1 cells exposed to **7** produced a single peak corresponding to the [M+H]^+^ion of **8**.

Analysis of 1D and 2D NMR data (gCOSY, gHSQC, gHMBC, ROESY, and TOCSY) allowed for structure determination of dihydroxypregnenolones **4** and **5**, corresponding to an expected 7α-hydroxylated product and an unexpected 7β-hydroxylated product (Figure 2B, Table S2). COSY interactions between the hydrogen at C-6 (δ_H_5.57, 5.28) and a hydroxyl-adjacent hydrogen (δ_H_3.8, 3.79) in **4** and **5**, respectively, revealed that both products were 7-hydroxylated. The C-7 hydroxyl stereochemistry was determined by comparing the isomers to reference NMR for 7α-hydroxypregnenolone (δ _C-7_65.20) and 7β-hydroxypregnenolone (δ _C-7_73.14) (Wang 2013, Schmitz 2014). Compound **4** (δ _C-7_65.95) is therefore 7α,17α-dihydroxypregnenolone while **5** (δ _C-7_73.99) is 7β,17α-dihydroxypregnenolone.

The CYP7B1 stable pool cells provided with alternative substrate **7** exhibited a single dihydroxyprogesterone product peak (Figure 3B). Dihydroxyprogesterones do not lose water as easily in ESI-MS, so the product peak was exclusively *m/z* = 347.222, corresponding to the [M+H]^+^ion. A single isomer **8** with this *m/z* eluted at 2.05 minutes, and was absent in the negative control lipids. 2.2 mg of **8** were eventually purified from tissue culture, with a yield of 6.9 μg mL^−1^.

Compound **8**, the single dihydroxyprogesterone product, was determined to be 11α,16α-dihydroxyprogesterone after analysis of 1D and 2D NMR data (Figure 2C, Table S2). No other masses that could correspond to the expected product **6** were detected. The newly hydroxylated carbon (δ_C_68.88, δ_H_3.95) was identified by locating its methylene neighbor (δ_C_50.89, δ_H_1.63,2.24), with which it has a strong COSY correlation. This neighboring carbon has an HMBC with hydrogens at C-17 (δ_H_2.58) and C-18 (δ_H_0.70). This placed the neighboring carbon at C-12, as the D-ring hydroxyl had already been assigned and other locations were too far from C-17 and C-18. Therefore the CYP7B1 hydroxylation was at C-11. The C-11 hydroxyl stereochemistry was found through ROESY interactions. The hydrogen at C-11 (δ_H_3.95) interacts with hydrogens at C-18 (δ_H_0.70) and C-19 (δ_H_1.35), indicating that it is above the steroid ring plane. The hydroxyl at C-11 must face the opposite direction, and thus is an 11α-hydroxyl.

The macromolecular modeling software Rosetta was used to model native substrate **1** and alternative substrates **3** and **7** binding to CYP7B1 to better understand how the unexpected products **5** and **8** could arise. Comparative modeling in Rosetta predicts protein structure based on solved structures of homologs, from which it generates candidate structures. CYP7A1, which is 41% sequence identical to CYP7B1 with 95% coverage, has been crystallized and served as a starting point for comparative modeling^18^. To account for induced fit in the active site, only CYP7A1 structures crystallized with substrates were used in templating (PDB IDs 3v8d and 3sn5). Steroids **1**, **3**, and **7** were each docked 10^4^ times into twenty holo-CYP7B1 structure models according to the RosettaLigand protocol^19^. Rosetta scores were used to select approximately seventy reasonable binding modes for each molecule. These modes were manually evaluated by proximity of the observed hydroxylation position to the ferryl oxygen and by similar orientation to steroids in homologous crystals. The resulting four binding modes were analyzed as the best hypotheses for how CYP7B1 catalyzes 7α-, 7β- and 11α-hydroxylation on the steroids studied (Figure 4).

**Figure 4.**
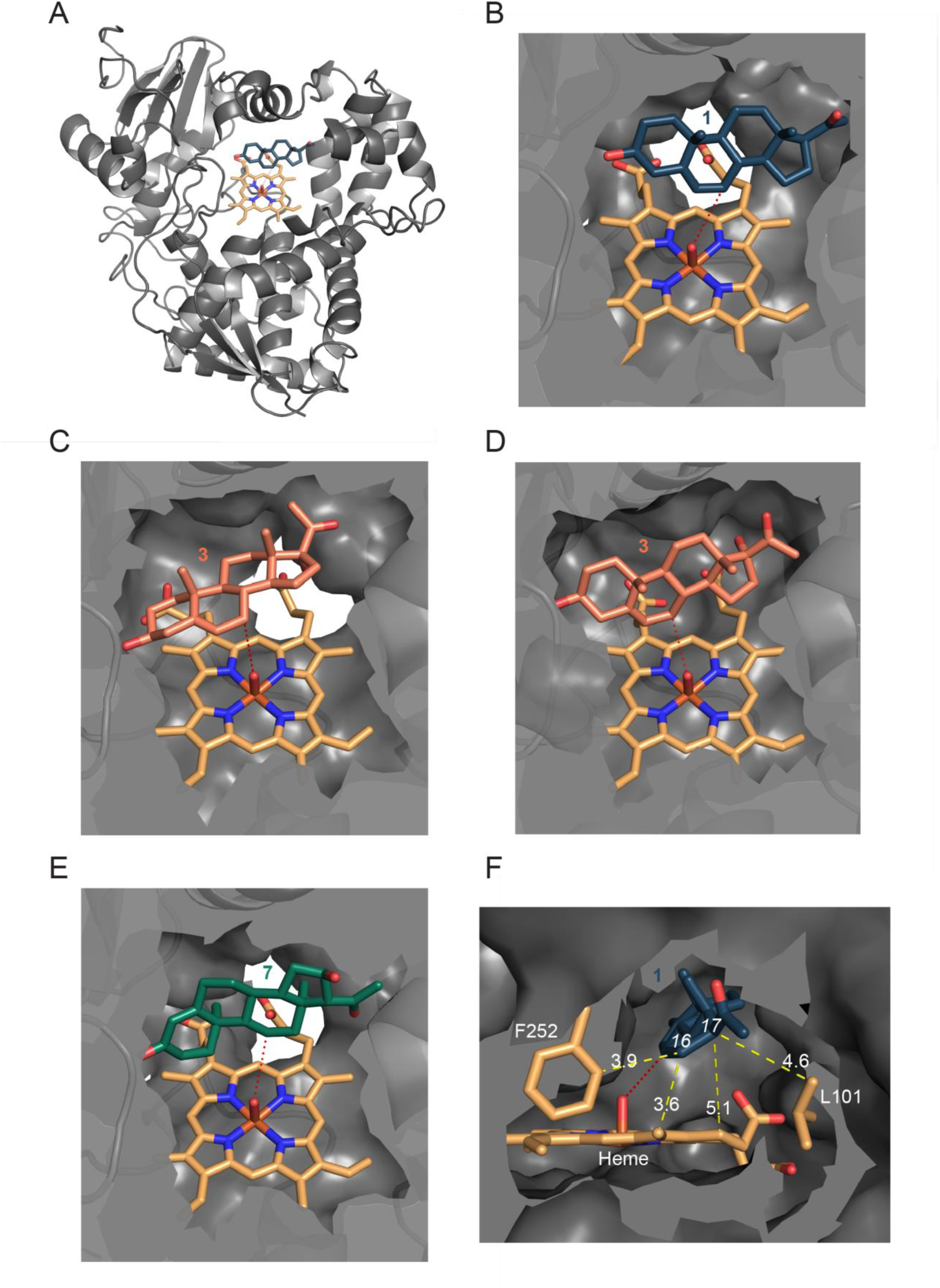
Predicted structures of the CYP7B1 active site with steroid substrates **1** (blue), **3** (orange), and **7** (green). A) Cartoon representation of holo-CYP7B1 with **1**. B-E) Structural models of CYP7B1 permitting 7α-hydroxylation of **1** (B) and **3** (C), 7β-hydroxylation of **3** (D), and 11α-hydroxylation of **7** (E). The red dashed line highlights the positions of the reactive ferryl oxygen and the observed hydroxylation site. F) Side view of native substrate **1** in active site. Yellow dashed lines indicate the proximity of steroid carbons C-16 and C-17 to the edge of the binding pocket. Distances are in Angstroms.

These structure models suggest that unexpected CYP7B1 activities on **3** and **7** arise from these substrates shifting out of a tight pocket and tilting relative to the reactive ferryl oxygen. Cytochrome P450s typically hydroxylate the carbon closest to the ferryl oxygen; the mechanism does not require any activation of the substrate, though the transient carbanion intermediate formed favors carbons adjacent to double bonds^20^. As C-7 is adjacent to a double bond in native substrate **1** and the alternative substrate **3**, it is a particularly good hydroxylation site, and thus observed in their products (Figure 4BC). However, **3**’s 17α-hydroxyl does not permit it to occupy the same binding position as **1** in our model. The hydroxyl forces **3** into a wider channel to the left in the figure, where the steroid can tilt to result 7β-hydroxylation (Figure 4D). **7**’s 16α-hydroxylation would also result in a steric clash, which implies it will also shift out of **1**’s binding pocket. The severity of this clash may drive **7** to rotate dramatically, resulting in the observed 11α-hydroxylation (Figure 4E). The side view of CYP7B1’s native substrate, pregnenolone (**1**), shows its tight fit in a hydrophobic binding pocket with little room beneath the D-ring (Figure 4F). This fit may hold the substrate steady to ensure hydroxylation exclusively at position 7α. Functional groups at C-16 and C-17, such as those in **3** and **7**, can disrupt this fit to cause the observed loss of reaction specificity.

We made the three steroids **4**, **5**, and **8** by expressing CYP7B1 in mammalian cells and providing it with non-native substrates **3** and **7**. **5** and **8** were the result of 7β- and 11α-hydroxylations, respectively, though CYP7B1 was thought to primarily perform 7α-hydroxylation. Docking substrates **3** and **7** into the enzyme active site suggests that these unexpected reactions arise from the substrates’ D-ring hydroxylations pushing them into a tilted orientation. This work represents a new approach in using engineered mammalian cells to produce novel steroids, and more broadly in using mammalian cells to produce novel small molecules. Our overall method can take advantage of inexpensive gene synthesis, diverse steroid-modifying enzymes, and mammalian cells’ unique properties to quickly create steroids that are compatible with direct biological testing.

Steroids **4** and **5** are novel molecules, but **8** has a straightforward organic synthesis. Nevertheless, **8**’s chemical properties are not published; no NMR structural data were released when it was initially isolated^21^. **4** and **5** would have been difficult to produce via established methods. Both have been tentatively identified in mixtures of microbial steroid hydroxylation products, but no NMR data is available^22,23^. Selective 7β-hydroxylation of pregnenolones is possible with organic methods, but syntheses that favor 7α-hydroxylation do not appear possible^24^. While microbial pregnenolone 7α-hydroxylation has been described, the organisms make a mixture of products and the enzymes have not been identified^25–27^. Thus our method for selective 7-hydroxylation fills a gap in steroid synthetic methods.

We observed unprecedented amounts of 11α-hydroxylated and 7β-hydroxylated products from CYP7B1. Prior to this work, CYP7B1 only performed 7α-hydroxylation, 6α-hydroxylation, or 7β-hydroxylation, depending on the substrate^28^. The lattermost product is minor, when observed at all; it comprised just 1% of the hydroxysteroid product from CYP7B1 action on DHEA^29^. In comparison, **5** was 36% of the total product isolated from CYP7B1 acting on **3**. 11α-hydroxylation by CYP7B1 had never been observed until we isolated **8**. These reactions demonstrate that even well-studied cytochrome P450s may process new substrates in unexpected ways; one cannot assume which isomers these enzymes produce based only on the available literature.

Our approach reveals that novel steroids can be generated by combining elements from the human metabolic repertoire. CYP7B1 is a human enzyme; its murine homolog is expressed primarily in the brain and at low levels in the liver^30^. Its 7α-hydroxylated steroid products are poorly understood, but improve memory in aged, memory-impaired mice^28,31^. While **7** is not present in the human body, **3** is an intermediate formed by CYP17A1 in the adrenal cortex and gonads^32^. However, human CYP17A1 preferentially processes **3** further into dehydroepiandrosterone^33^. Though the enzyme dissociates from the steroid after hydroxylation, relatively little **3** can escape into other tissues before CYP17A1 acts again^34^. While CYP7B1 likely does not encounter **3** in nature, in our system the enzyme is capable of processing this substrate into novel products **4** and **5**.

Here, we engineered mammalian cells to produce previously uncharacterized drug-like small molecules. Numerous protein drugs are already made in mammalian cells, typically due to species-specific post-translational modifications such as N-linked glycosylation. Mammalian cells and mammal-based cell-free systems have been engineered to make steroids, but these were only used to study the biosynthetic pathways of known molecules^11,35^. Engineered steroid biosynthesis, even in microbes, has only been used to make known steroids; novel steroids are detected spontaneously from unmodified cells^9,12,25,36^. This work therefore opens the door both towards using mammalian cells for other small molecule products, and also towards directed biosynthesis of novel steroids.

Many novel molecules could be made with engineered cells expressing steroid-modifying enzymes, making more steroids accessible for biological testing. The decreasing cost of gene synthesis, the high frequency of mammalian cell co-transfection when plasmids are simply mixed together, and the high activity of steroids in biological assays could enable a means of identifying useful compounds. Our *in vivo* system could be modified to access all parts of the steroid ring system, unlike current organic synthesis methods. Specifically, we envision that libraries of steroids could be made by combinatorial transfection of mammalian cells, followed for example by direct biological testing in high-content cell-based assays to find new activities^37^.

## Methods

### Plasmids

The CYP7B1-containing plasmid had a pcDNA3.1 backbone, a CMV promoter driving intron-free CYP7B1 and puromyin resistance. The negative control plasmid was identical, but contained GFP in place of CYP7B1. A list of all steroid-modifying enzymes used in similar plasmids can be found in Table S1. Plasmids were constructed using Gibson assembly to insert IDT gBlocks containing the gene of interest into PCR-amplified plasmid backbones, which were then transformed into chemically-competent *Escherichia coli* K12. Plasmids were prepared using the Qiagen PlasmidPlus MidiPrep or MaxiPrep kit.

### Protein Expression in Mammalian Cells

HEK293 cells (ATCC) were grown in DMEM (Invitrogen) with 10% FBS (Gibco) and penicillin-streptomycin (Gibco). Cells were transfected with 31.5 μg of plasmid DNA per well in 6-well plates using the Lipofectamine 3000 kit (Thermo Fisher). Media was replaced after six hours with fresh media containing 40 μM steroid substrate. Cells were incubated for 48 hours, after which the negative controls were checked for fluorescence, indicating acceptable transfection efficiency. Spent media and cells were mixed by scraping, and the liquid was frozen for storage. Stable pools of HEK293 cells expressing steroid-modifying genes began with transfection of ∼800,000 cells using the lipofectamine 3000 kit, as above. 48 hours after transfection, cells were selected via 1.5 μg mL^−1^ puromycin (Sigma). After eighteen days, antibiotic was decreased to 0.5 μg mL^−1^puromycin. To modify steroids, stable pool cells were plated at 7,000 cells per cm^2^ with 40 μM of the desired substrate. The cells were incubated for eight days, after which adhered cells were scraped to mix with the media and frozen for storage.

### Steroid Product Isolation

Liquid-liquid separation with a 3:10 methanol: methyl-tert-butyl ether v/v organic phase removed lipids from the media^14^. The 2 mL of media was extracted first with 6.5 mL and then with 3 mL of solvent, vortexing for three minutes each time. The upper layers were removed, combined, dried down, and resuspended in 100 μL of methanol. Lipids were analyzed with an Agilent 1200 series HPLC system and 6530 qTOF mass spectrometer. Steroids were separated with a Thermo Scientific Hypersil GOLD C18 column (1.9 μm, 50 x 2.1 mm) in an acetonitrile gradient in water with 0.1% formic acid v/v^15^. Known products were compared to standards for quantification. Please refer to the supporting information for HPLC gradients, steroid sources, and chromatograms of steroid products with standards.

### Unknown Steroid Purification and NMR

Twenty-two T150 flasks of the CYP7B1-expressing cells were used to process substrate **3**, yielding 440 mL of media. Sixteen similar flasks were used to process substrate **7**, resulting in 320 mL of media. Liquid-liquid extraction was performed with the same proportions as above, with the addition of a sodium sulfate drying step. The product mixtures from **3** and **7** were resuspended in 3 and 5 mL of methanol, respectively. An Agilent 1200 semi-preparative HPLC system with a Phenomenex Luna C18 column (5 μm, 250 x 10 mm) was used to purify the steroid products **4**, **5**, and **8**. This resulted in 0.7 mg of **4**, 0.4 mg of **5**, and 2.2 mg of **8**. NMR spectra were obtained in CD_3_OD with a Bruker AVANCE 500 MHz spectrometer equipped with a ^1^H{^13^C/^15^N} cryoprobe and a Bruker AVANCE 500 MHz spectrometer equipped with a ^13^C/^15^N{^1^H} cryoprobe. ^13^C and ^1^H shifts and assignments are in Table S2. ^13^C, ^1^H, HSQC, HMBC, COSY, ROESY, and TOCSY spectra, along with HPLC methods, can be found in the supporting information.

### Rosetta Steroid Docking Models

We generated comparative models for CYP7B1 binding steroids based on crystallized enzymes with Rosetta^38^. The algorithm selects for structures with optimal physical interactions, solvent accessible surface area, and bond angles by assigning structures a Rosetta score; lower Rosetta Energy Unit (REU) values suggest more energetically favorable conformations^38^. The two solved structures within 30% sequence identity and 10^−4^ BLAST confidence, and bound to chemically similar substrates, were PDB IDs 3sn5 and 3v8d^18^. We threaded the sequence of CYP7B1 onto the coordinates of those structures and combined fragments of each to create ten thousand hybrid models of CYP7B1^19,38^. The heme was inserted according to the original crystals’ bond angles and obabel-derived partial charges; this was followed by all-atom optimization^39^. ∼2500 reasonable structures were identified by their Rosetta score falling within fifty REU of the lowest-scoring structure, and were clustered at 3 Å RMSD. An ensemble of the twenty best-scoring CYP7B1 conformations were used in docking **1**, **3**, and **7**, generating ten thousand binding predictions for each steroid. Typical parameters for these RosettaLigand docking runs were used: the steroid was randomly placed within 5 Å of the binding pocket center, coarsely fit inside, and accommodated by sidechains within 5 Å^19^. Energetically feasible binding modes were selected by an overall Rosetta score within 65 REU of the best mode by overall score, and a ΔG-binding analog score within 4 REU of the best mode by ΔG-binding score. Binding modes were only considered if at least one atom of the steroid was within 4 Å of the ferryl oxygen; this was true for approximately four in five of the generated structures. Selected structures were clustered at 3 Å RMSD between substrate atoms, and the cluster representative was chosen by the lowest ΔG-binding analog. This resulted in approximately seventy structures per steroid, which were manually evaluated. We selected for conformations that placed the hydroxylation site within 3.5 Å of the ferryl oxygen, and against conformations that would require unlikely substrate trajectories into the active site. Thus we produced a binding mode for each hydroxylation mode for each substrate, for a total of four models. Please refer to the supporting information for final and intermediate structure model files, as well as clustering details.

## Acknowledgements

This work was supported by the Defense Advanced Research Projects Agency HR0011-12-C-0061 and the National Institute of Health P50 GM107618 (Laboratory of Systems Pharmacology) and R01 GM086258 (JC). We are grateful to the ICCB-Longwood Analytical Chemistry Core facility at Harvard Medical School for use of their mass spectrometer. We would also like to thank S. Trauger, K. Chatman, and G. Byrd at the Harvard Small Molecule Mass Spectrometry Core.

## Supporting Information Available

Yield tables for all enzymes, descriptions of conditions to improve yield, supplementary methods, and NMR shifts, assignments, and spectra (PDF). CYP7B1 substrate binding predictions from Rosetta (ZIP). Intermediate structures used in CYP7B1 modeling (ZIP). This material is available free of charge via the Internet at http://pubs.acs.org.

The authors declare no competing financial interest.

